# Visual Function Correlates More Strongly with Glial Coverage than Axon Count Across Multiple Mouse Strains

**DOI:** 10.64898/2026.03.23.713746

**Authors:** Benton Chuter, William White, T.J. Hollingsworth, XiangDi Wang, Linan Guan, Min Young Kim, Monica M. Jablonski

**Affiliations:** Department of Ophthalmology, Hamilton Eye Institute, The University of Tennessee Health Science Center, Memphis, TN, United States; Department of Ophthalmology, The First Affiliated Hospital of Harbin Medical University, Harbin, Heilongjiang, China

**Keywords:** Glaucoma, Optic nerve, Morphometrics, Retinal ganglion cells, Structure-function, Mouse model

## Abstract

**Objective:** To determine whether non-axon optic nerve morphometric features correlate with clinical visual function as strongly as the traditional axon count gold standard.

**Design:** Cross-sectional histological analysis with longitudinal clinical correlation.

**Subjects:** Eighteen mice from three strains: C57BL/6J (n=6), BXD51 (n=6), and DBA/2J (n=6).

**Methods:** Left eye (OS) optic nerves from mice euthanized at 12 months of age were resin-embedded and stained with p-phenylenediamine. Bright-field cross-sectional images were segmented using an AxonDeepSeg-based workflow to generate axon, myelin, whole nerve, and glial coverage masks for morphometric quantification. Seven morphometrics were extracted: axon count (nAx), axon density (AxDen), glial coverage area ratio (GliaR), mean solidity (Sol), mean axon diameter (AxDiam), mean myelin area (MyArea), and mean axon-myelin area (AxMyArea). Morphometrics were correlated with longitudinal clinical data collected at 1, 3, 6, 9, and 12 months, including visual acuity (VA), contrast threshold, intraocular pressure (IOP), and pattern electroretinography P50 and N95 amplitudes (PERG P50 and N95).

**Main Outcome Measures:** Pearson correlation coefficients were used to assess associations between morphometric features and clinical measures, and Fisher z-transformed meta-analytic correlations were used to aggregate these associations across ages.

**Results:** VA and contrast threshold demonstrated strong correlations with GliaR that matched or exceeded nAx. Meta-analysis across ages revealed GliaR correlated with VA (r = -0.84, *p* = 4.49 × 10^-21^) and contrast threshold (r = 0.86, *p*=7.55 × 10-^23^), comparable to nAx correlations with VA (r = 0.80, *p*=8.13×10^-17^) and contrast threshold (r = -0.80, p= 1.74×10^-16^). Structure-function relationships shifted with age: at 6 months, GliaR had the strongest correlation with contrast threshold (r = 0.96), while at 12 months, AxDiam became the dominant correlate of both VA (r = 0.77) and contrast threshhold (r = -0.74). IOP, PERG P50, and PERG N95 exhibited weak correlations with all morphometrics (|r| < 0.27).

**Conclusions:** Non-axon morphometrics, particularly glial coverage area ratio, correlate with visual function as strongly as traditional axon count. Automated optic nerve assessment should incorporate glial and other non-axon features. Further, stage-aware biomarker selection may better capture structure-function relationships in glaucoma.

## INTRODUCTION

Glaucoma remains the leading cause of irreversible blindness worldwide (Tham et al., 2014), characterized by progressive degeneration of retinal ganglion cells (RGCs) and their axons (Howell et al., 2007). Axon counting has long served as the historic gold standard for quantifying optic nerve damage and validating structural biomarkers in glaucoma research (Reynaud et al., 2012; Cull et al., 2012). This metric provides the reference standard for assessing disease severity, evaluating neuroprotective interventions, and validating structure-function models that link anatomical changes to visual outcomes (Harwerth et al., 2010). However, the optic nerve comprises multiple features relevant to optic nerve health beyond axons alone. Astrocytes, the predominant glial cell type in the optic nerve head, undergo pronounced reactive changes during glaucomatous neurodegeneration (Hernandez, 2000), which include gliosis characterized by increased glial fibrillary acidic protein expression, hypertrophy, and migration to sites of damage (Tezel et al., 2001). These glial coverage changes, including gliosis and glial scar formation, may precede, accompany, and/or persist beyond acute axonal loss (Dai et al., 2012). Despite growing recognition of glial involvement in glaucoma pathophysiology, quantitative assessment of non-axon microstructural features and their relationship to visual function remains limited.

The advent of deep learning-based segmentation tools has enabled automated, high-throughput quantification of optic nerve morphometry. Tools, such as AxonDeepSeg, can segment axons and myelin sheaths from microscopy images with high accuracy across species (Zaimi et al., 2018). These tools have primarily been applied to extract axon counts and myelin metrics, rather than quantify glial coverage area ratios and other non-axon features that may carry independent information about tissue health. Because axonal loss is accompanied by glial expansion and structural remodeling, quantifying these features may provide additional insight into optic nerve health. Furthermore, although structure-function relationships in glaucoma may vary across disease stages, stage-dependent correlations between morphometric features and visual function have not been established in mouse models (Medeiros et al., 2012). Understanding how these relationships evolve with age and disease progression could inform the selection of stage-appropriate biomarkers for monitoring and intervention studies.

In this study, we investigated whether non-axon morphometric features, particularly glial coverage area ratio, correlate with visual function measures as strongly as axon count. We further assessed age-dependent changes in structure-function relationships and compared these associations across three mouse strains representing a range of optic nerve phenotypes: C57BL/6J (B6; control), BXD51, and DBA/2J (D2; glaucoma model) (Schlamp et al., 2006; Geisert & Williams, 2020). We hypothesized that glial coverage area ratio would correlate with visual acuity (VA) and contrast threshold at a level comparable to axon count. We show that these relationships hold across strains, reflecting the broad relevance of glial changes in glaucomatous neurodegeneration.

## METHODS

### Animals and Study Design

Three mouse strains representing distinct glaucoma phenotypes were studied: C57BL/6J (B6, control), BXD51 (recombinant inbred, intermediate glaucoma phenotype), and DBA/2J (D2, severe inherited glaucoma). Six mice per strain (n=18 total) were included in the final analysis. Each strain included balanced sex representation (3 males, 3 females per strain). All mice underwent longitudinal clinical assessments at 1, 3, 6, 9, and 12 months of age, followed by sacrifice at 12 months for optic nerve histological assessment. Left eyes (OS) were used for morphometric-clinical correlation analyses to maintain independence between observations. All procedures were approved by the University of Tennessee Health Science Center (UTHSC) IACUC (Protocol #24-0543) and performed in accordance with the Association for Research in Vision and Ophthalmology Statement for the Use of Animals in Ophthalmic and Vision Research. Mice were housed at the UTHSC lab animal care unit under a 12-hour light, 12-hour dark illumination cycle, with food and water freely available.

### Clinical Visual Function Assessments

#### Visual Acuity

Spatial frequency thresholds were measured using the optokinetic tracking response using the OptoDrum system (Striatech, Tübingen, Germany). Mice were placed on an elevated platform surrounded by rotating vertical sine-wave gratings. The spatial frequency of the grating was increased in a staircase procedure until the reflexive head-tracking response was no longer elicited, defining the visual acuity threshold in cycles per degree.

#### Contrast Threshold

Contrast thresholds were assessed as a measure of visual function using the same optokinetic apparatus. Grating contrast was reduced at a fixed spatial frequency until the tracking response was extinguished, and the minimum contrast required to elicit tracking was recorded as the contrast threshold.

#### Intraocular Pressure

Intraocular pressure (IOP) was measured using a TonoLab tonometer (iCare USA, Raleigh, NC). Measurements were obtained in mice lightly sedated with isoflurane, during the same time window each testing day to minimize diurnal variation.

#### Pattern Electroretinography

Pattern electroretinography was performed using the Celeris system (Diagnosis LLC, Lowell, MA). The amplitudes of the P50 and N95 waveforms were measured, reflecting pre-ganglionic retinal input and RGC function, respectively. Amplitudes were measured from baseline to peak for P50 and from peak to negative trough for N95.

### Optic Nerve Histology

At 12 months, mice were euthanized, and eyes with attached optic nerves were enucleated and immersion-fixed in a mixture of 2% paraformaldehyde and 2% glutaraldehyde in phosphate buffer. Optic nerves were dissected posterior to the globe, fixed, dehydrated, and embedded as previously described (Stiemke et al., 2020). One-micron thick cross-sections were cut and stained with p-phenylenediamine (PPD) to visualize myelin sheaths and degenerating axons under bright-field microscopy using a ZEISS Axio observer 7 inverted microscope (Carl Zeiss, Jena, Germany). Images were acquired as multiple individual fields at high magnification using a 63x objective, which were digitally stitched using the image tiling feature to generate a composite image of the entire optic nerve cross-section.

### Automated Morphometric Quantification

Optic nerve cross-section images were analyzed using a custom pipeline built on AxonDeepSeg, a validated deep learning framework for axon and myelin segmentation (Zaimi et al., 2018; Chuter et al., 2026, Preprint). The pipeline calibrated pixel dimensions, then performed convolutional neural network-based segmentation of individual axons and their myelin sheaths. A morphology-based contour extractor merged these axon and myelin masks into a nerve core and applied basic morphology and gradient-plus-spline smoothing to generate whole-nerve contour masks. These contour masks yielded a total cross-sectional area, and glial coverage regions were identified by subtracting axon-myelin masks from whole-nerve contour masks. All segmentation outputs underwent visual quality control review. Axon counts derived from light microscopy and deep learning segmentation are expected to underestimate total axon numbers compared to transmission electron microscopy, as smaller unmyelinated axons fall below the resolution threshold of the imaging system (Zaimi et al., 2018).

Seven morphometric features were extracted from each optic nerve: (1) axon count (nAx): total number of segmented axons per cross-section; (2) axon density (AxDen): number of axons per pixel of cross-sectional area; (3) glial coverage area ratio (GliaR): ratio of glial coverage area to total nerve cross-sectional area; (4) mean solidity (Sol): average axon solidity indicating axon shape regularity; (5) mean axon diameter (AxDiam): average equivalent circular diameter of axons in micrometers; (6) mean myelin area (MyArea): average myelin sheath cross-sectional area; and (7) mean axon-myelin area (AxMyArea): average total fiber cross-sectional area. Morphometric values were reported as mean ± SD.

### Statistical Analysis

Pearson correlation coefficients were calculated between each morphometric feature and each clinical measure at each age. To obtain aggregate effect estimates across ages, individual correlation coefficients were Fisher z-transformed, pooled as weighted means, and back-transformed to the correlation scale. Ninety-five percent confidence intervals and p-values were derived from the meta-analytic z-statistics. Multivariate analysis of variance (MANOVA) tested the overall effect of strain on the seven-morphometric profile. One-way ANOVA with Tukey HSD post-hoc comparisons assessed strain differences for individual morphometrics. Two-way ANOVA assessed main effects of strain and sex, and their interaction; within-strain sex comparisons used unpaired t-tests. Principal component analysis was applied to standardized morphometric data to visualize strain separation. Statistical significance was defined as two-tailed *p*<.05.

## RESULTS

### Morphometric and Clinical Profiles Differ by Strain

Morphometric analysis of optic nerve cross-sections from 12-month-old mice using automated morphometric quantification (**Supplementary Figure S1**) revealed pronounced differences among strains (**Figure 1, Table 1**). Axon counts were highest in B6 mice (33,224 ± 5,660 axons), intermediate in BXD51 mice (28,438 ± 2,763 axons), and lowest in D2 mice (11,272 ± 11,250 axons). Glial coverage area ratio followed an inverse pattern: lowest in B6 (0.18 ± 0.03), intermediate in BXD51 (0.21 ± 0.03), and highest in D2 (0.42 ± 0.09), consistent with increased gliosis accompanying axon loss.

**Table 1.** Optic Nerve Morphometric Profiles by Strain at 12 Months.

| Morphometric parameter | C57BL/6J (B6, n=6) | BXD51 (n=6) | DBA/2J (D2, n=6) | *p*-value |
| --- | --- | --- | --- | --- |
| Axon count | 33,224 ± 5,660 | 28,438 ± 2,763 | 11,272 ± 11,250 | .0003 |
| Axon density (x10<sup>-4</sup>/px) | 4.58 ± 0.49 | 4.56 ± 0.50 | 1.52 ± 1.44 | .0003 |
| Glial coverage area ratio | 0.18 ± 0.03 | 0.21 ± 0.03 | 0.42 ± 0.09 | <.00001 |
| Mean solidity | 0.94 ± 0.003 | 0.94 ± 0.003 | 0.93 ± 0.009 | .0003 |
| Mean axon diameter (um) | 1.71 ± 0.12 | 1.56 ± 0.06 | 1.47 ± 0.05 | .0003 |
| Mean myelin area (um<sup>2</sup>) | 5.35 ± 0.63 | 5.12 ± 0.39 | 3.91 ± 0.81 | .003 |
| Mean axon-myelin area (um<sup>2</sup>) | 8.37 ± 1.05 | 7.66 ± 0.45 | 6.29 ± 0.76 | .003 |

Morphometric characteristics of optic nerve cross-sections are presented as mean ± SD by strain. *P*-values from one-way ANOVA assess whether the means across the three strains differ significantly for each morphometric.

**Figure 1.**
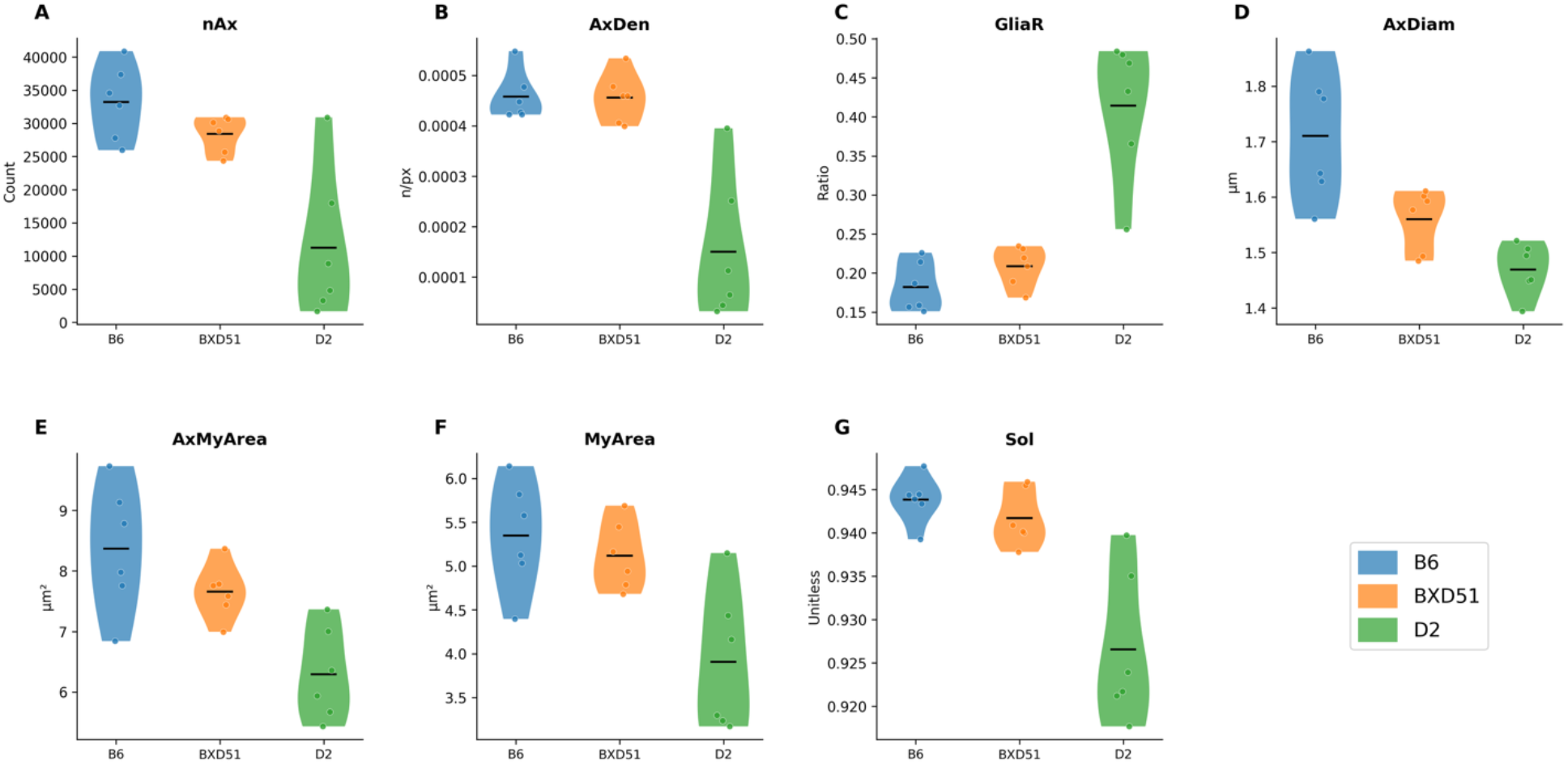
Optic nerve morphometric profiles across mouse strains with distinct glaucoma phenotypes. Seven morphometric features were measured at 12 months of age in C57BL/6J (B6), BXD51, and DBA/2J (D2) mice (n = 6 per strain). Quantified parameters were **(A)** Axon count, **(B)** axon density, **(C)** glia coverage area ratio (GliaR), **(D)** mean solidity, **(E)** mean axon diameter, **(F)** mean myelin area, **(G)** mean axon-myelin area. Horizontal lines indicate group medians.

Clinical visual function measures also differed significantly among strains (**Supplementary Table S1, Supplementary Figure S2**). Contrast threshold was lowest in B6 mice, intermediate in BXD51, and highest in D2 mice (B6 vs D2: mean difference = 84.97, *p*<.001; BXD51 vs D2: mean difference = 54.7, *p*<.001; B6 vs BXD51: mean difference = 30.27, *p*<.001). Visual acuity was highest in B6 with BXD51 exhibiting comparable acuities that exceeded D2 values (B6 vs D2: mean difference = 0.30 cycles/degree, *p*<.001; BXD51 vs D2: mean difference = 0.25 cycles/degree, *p*<.001; B6 vs BXD51: *p*=.98, not significant).

### Sex Effects on Morphometric Parameters

Sex-stratified analysis revealed significant strain × sex interactions for mean axon diameter (*p*=.011) and mean axon+myelin area (*p*=.035), indicating that sex effects varied by strain. In B6 controls, males exhibited larger mean axon diameters than females (1.81 ± 0.05 vs 1.61 ± 0.04 μm; *p*=0.006) and lower glial coverage area ratios (0.16 ± 0.004 vs 0.21 ± 0.020; *p*=.010). Neither BXD51 nor D2 had significant sex differences for any morphometric parameter (*p*>.05; **Supplementary Table S2, Supplementary Figure S3**).

### Glial coverage Area Ratio Correlates with Visual Function as Strongly as Axon Count

Fisher z-transformed meta-analysis across all ages revealed that glial coverage area ratio (GliaR) correlated with VA and contrast threshold as strongly as traditional axon count (**Table 2**). Correlations with visual acuity were |r| = 0.84 (95% CI: -0.90 to –0.75, *p*=4.49 × 10^-21^) for GliaR and r = 0.80 (95% CI: 0.68 to 0.87, *p*=8.13 × 10^-17^) for axon count. For contrast threshold, the strongest correlation was observed for GliaR (r = 0.86, 95% CI: 0.78 to 0.92, *p*=7.55 × 10^-23^), exceeding that of axon count (|r| = 0.80, 95% CI: -0.87 to –0.68, *p*=1.74 × 10^-16^).

**Table 2.**
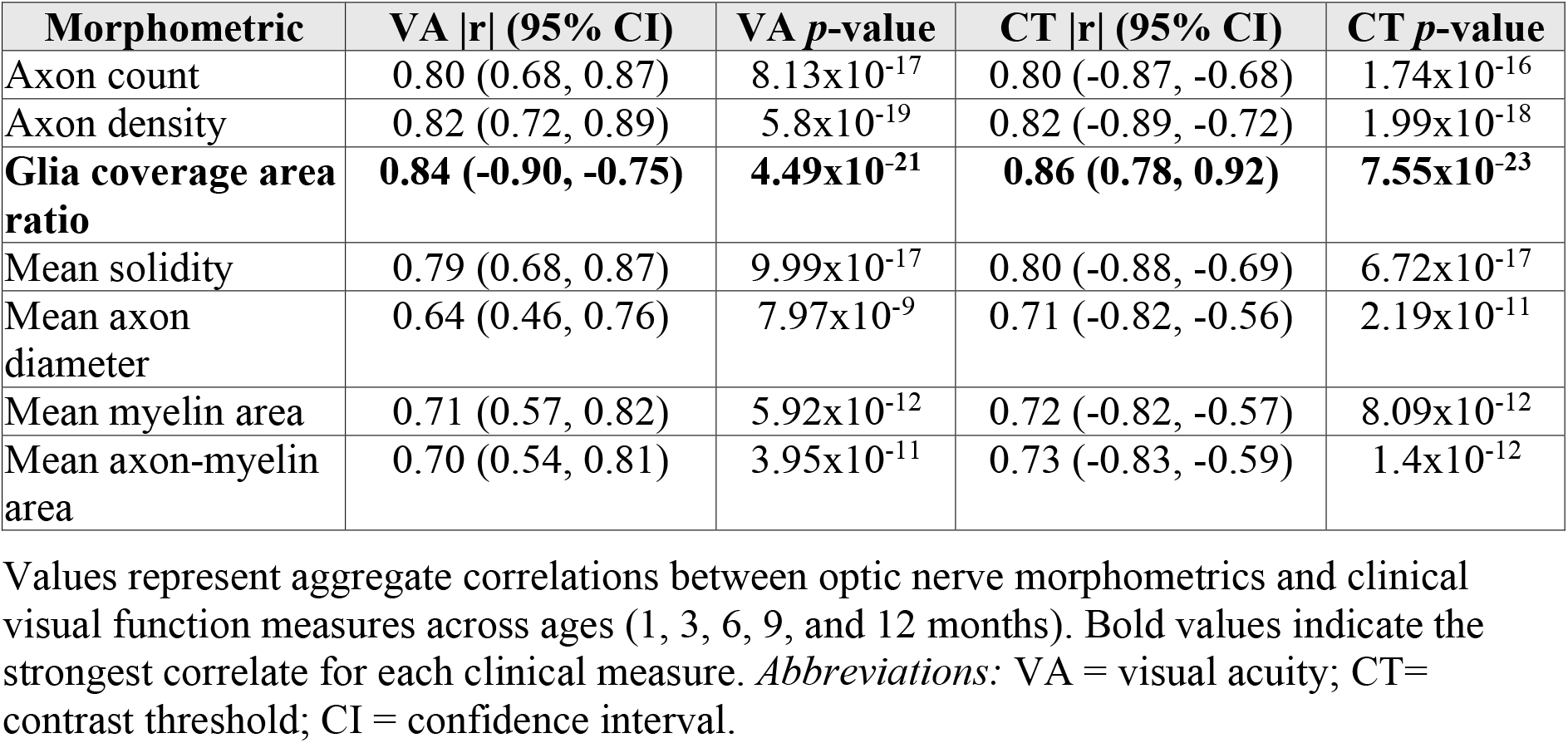
Fisher z-transformed Meta-Analytic Correlations Between Morphometrics and Clinical Measures.

In contrast, correlations between IOP and morphometric features were weak and not statistically significant (|r| < 0.23 for all, *p*>.05). Pattern electroretinography P50 amplitude (PERG P50) also exhibited weak, non-significant correlations with all morphometrics (|r| < 0.24, *p*>.05). Correlations between PERG N95 and morphometrics were also weak, with axon diameter approaching significance (r = 0.27, *p*=.055) but all other correlations non-significant (|r| ≤ 0.14).

### Structure-Function Relationships Shift with Age

Age-stratified correlation analysis revealed that the morphometric feature most strongly associated with visual function changed across the disease course (**Table 3, Figures 2 and 3**). Absolute correlation magnitudes organized by clinical measure (**Figure 2**) demonstrated that VA and contrast threshold consistently maintained strong associations with multiple morphometrics across ages, whereas IOP, PERG P50, and PERG N95 had weaker and more variable correlations. Signed correlation heatmaps (**Figure 3**) revealed the polarity of these relationships: axon-related metrics (nAx, AxDen, Sol, AxDiam, MyArea, AxMyArea) correlated positively with VA and negatively with contrast threshold, whereas GliaR exhibited consistent correlated negatively with VA and positively with contrast threshold.

**Table 3.**
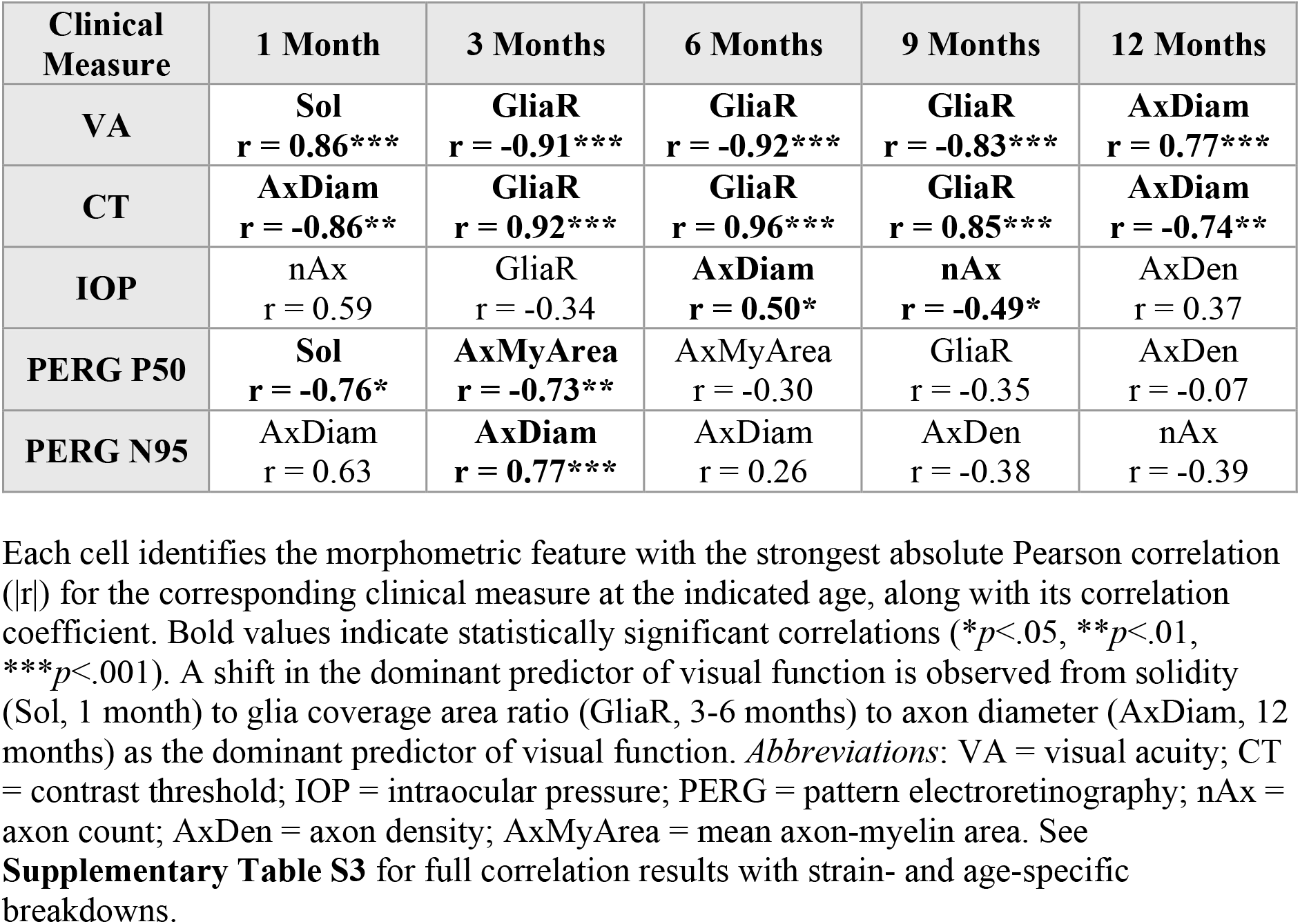
Top Morphometric Predictor of Each Clinical Measure by Age.

**Figure 2.**
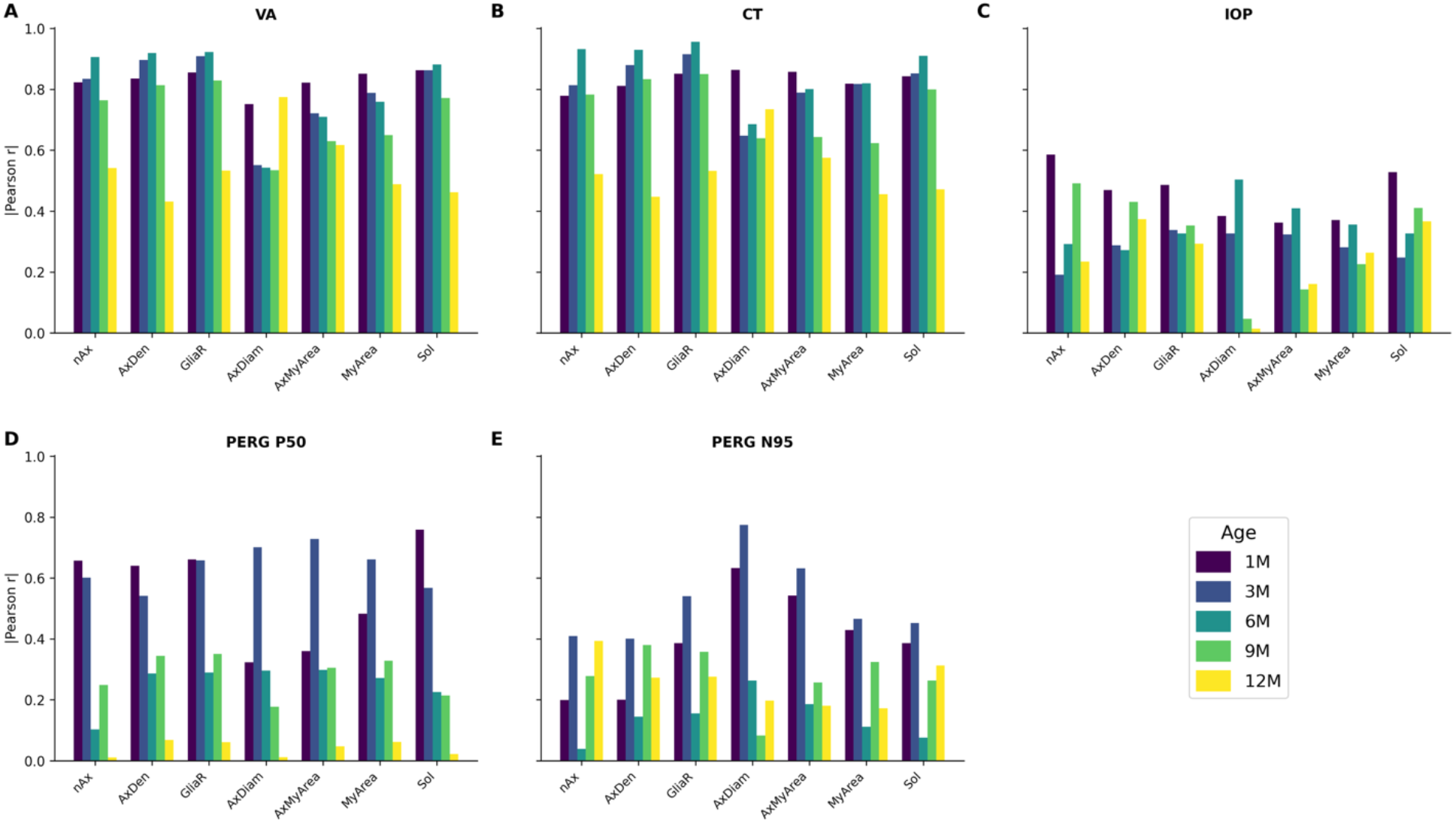
Absolute Pearson correlations between optic nerve morphometrics and clinical measures across ages. Absolute correlation coefficients (|r|) are shown for each morphometric feature relative to **(A)** visual acuity (VA), **(B)** contrast threshold (CT), **(C)** intraocular pressure (IOP), and **(D)** pattern electroretinography P50 (PERG P50) and **(E)** PERG N95 amplitudes at 1, 3, 6, 9, and 12 months (labeled 1M, 3M, 6M, 9M, and 12M in the legend). Morphometric features include axon count (nAx), axon density (AxDen), glial coverage area ratio (GliaR), mean axon diameter (AxDiam), mean axon–myelin area (AxMyArea), mean myelin area (MyArea), and mean solidity (Sol).

**Figure 3.**
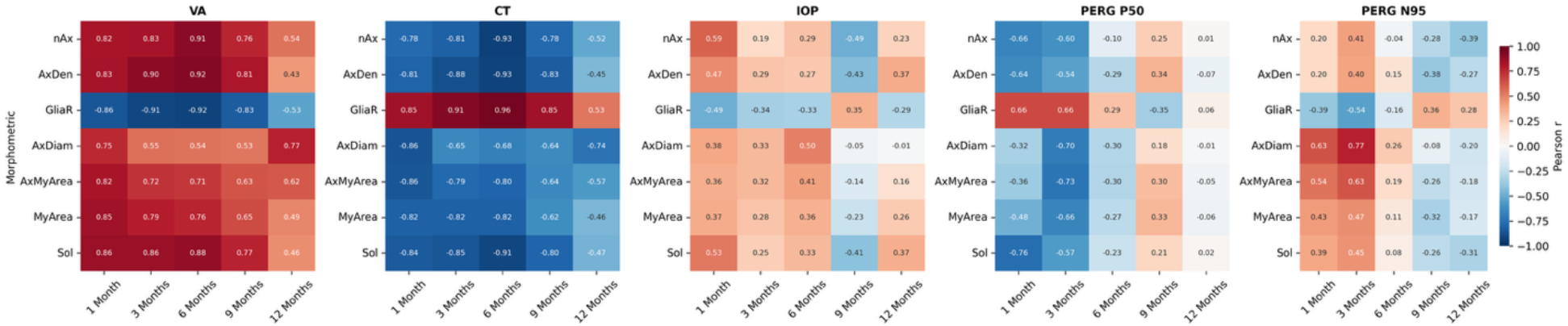
Signed Pearson correlation heatmaps reveal age-dependent direction and magnitude of morphometric-clinical associations. Heatmaps display Pearson correlation coefficients (r) between morphometric features (nAx = axon count; AxDen = axon density; GliaR = glial coverage area ratio; AxDiam = mean axon diameter; AxMyArea = mean axon-myelin area; MyArea = mean myelin area; Sol = mean solidity) and clinical measures (VA = visual acuity; CT = contrast threshold; IOP = intraocular pressure; PERG P50 and N95 = pattern electroretinography P50 and N95) at each age (1, 3, 6, 9, and 12 months). Negative correlations between GliaR and VA are observed across multiple ages, and the strongest morphometric correlates vary with age. Red indicates positive correlations, and blue indicates negative correlations, with color intensity reflecting correlation magnitude.

At early timepoints (1 month), the strongest correlation with VA was observed for mean solidity (r = 0.86, *p*<.001), whereas axon diameter was most strongly correlated with contrast threshold (r = -0.86, *p<*0.01; **Table 3**). During mid-disease (3-6 months), the GliaR emerged as the dominant correlate of VA. At 6 months, the strongest correlation with contrast threshold was observed for GliaR (r = 0.96, *p*=2.82 × 10^-8^). At the late timepoint (12 months), the strongest structure-function correlations with both VA (r = 0.77, *p*<0.001) and contrast threshold (r = -0.74, *p<*0.01) were observed for axon diameter.

### Morphometric-Clinical Relationships Are Consistent Across Strains

Principal component analysis of the seven morphometric features revealed clear strain separation (**Figure 4**). B6 mice clustered in the region of high axonal health, D2 mice in the region of low axonal health and high glial coverage ratio, and BXD51 mice occupied an intermediate position between these extremes. Notably, despite their distinct morphometric profiles, all three strains contributed to the overall structure-function correlations. The strong correlations between GliaR and visual function were not driven solely by D2 mice with severe pathology; rather, the relationship held across the continuum of disease severity represented by the three strains.

**Figure 4.**
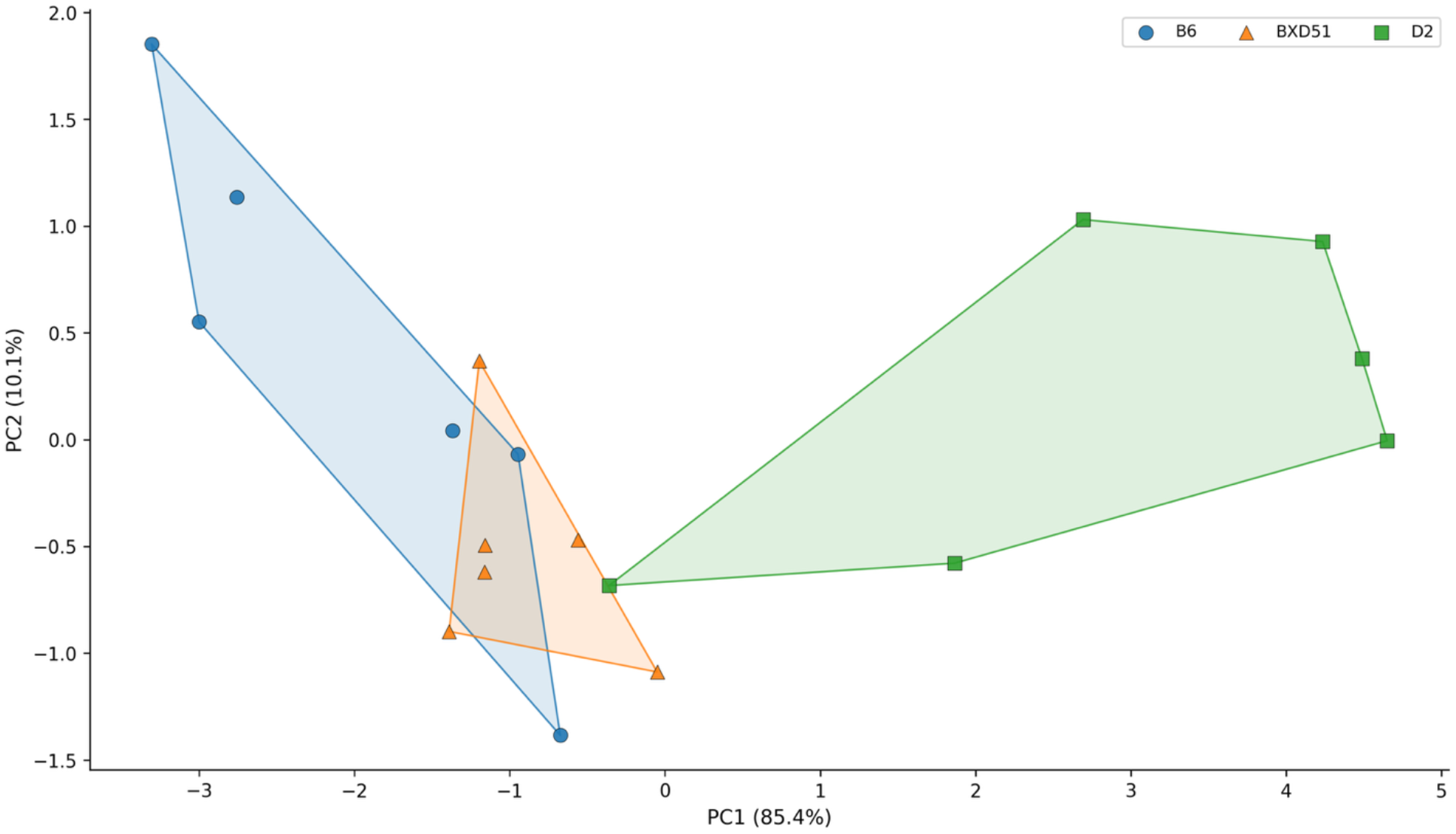
Principal component analysis of optic nerve morphometrics across mouse strains. Individual optic nerves from C57BL/6J (B6), BXD51, and DBA/2J (D2) mice are projected onto the first two principal components (PC) derived from seven standardized morphometric features (PC1, 85.4% variance; PC2, 10.1% variance; n = 6 for each mice strain). PC1 represents a composite axis of optic nerve health, with negative values associated with higher axon count, axon density, solidity, and myelin area, and positive values associated with higher glial coverage ratio. B6 nerves cluster at negative PC1 values, D2 nerves cluster at positive PC1 values, and BXD51 nerves fall in intermediate position along this axis, consistent with a continuum of glaucomatous damage across strains.

## DISCUSSION

Mouse models spanning a range of disease severities can test whether morphometric biomarkers generalize across the glaucoma spectrum. The D2 strain develops pigmentary dispersion glaucoma with progressive optic nerve degeneration that becomes apparent between 8-9 months and severe by 12 months of age (Libby et al., 2005; Schlamp et al., 2006). B6 mice serve as controls with more preserved optic nerve structure. BXD recombinant inbred strains between D2 and B6 exhibit variable phenotypes that may represent intermediate disease states (Geisert & Williams, 2020). Comparing structure-function relationships across strains with distinct pathological profiles help establish whether specific morphometric features consistently predict visual outcomes regardless of genetic background or disease severity.

In the present study, visual function was assessed using both VA and contrast threshold. Higher threshold values indicate poorer contrast sensitivity, reflecting a greater contrast requirement to elicit a visual response (Douglas et al., 2005). Additionally, D2 mice exhibit progressive RGC dysfunction and visual deficits, supporting the interpretation of elevated contrast threshold as a marker of impaired visual function in glaucomatous disease (Porciatti et al., 2007).

This study demonstrates that glial coverage area ratio, a metric reflecting glial and extracellular matrix content in the optic nerve, correlates with visual function measures as strongly as traditional axon count in mouse models spanning the glaucoma severity spectrum. The finding that GliaR showed the highest correlation with both VA (r = -0.84) and contrast threshold (r = 0.86) across ages suggests that non-axon compartments carry independent and clinically relevant information about optic nerve health. These results challenge the long-standing view that axon count alone is sufficient for characterizing structure-function relationships in glaucomatous optic neuropathy.

The magnitude of correlations observed in this study compares favorably with structure-function relationships reported in larger mammalian models. Harwerth et al. reported correlations between visual sensitivity and RGC counts ranging from r = 0.65 to 0.85 in primate glaucoma models (Harwerth et al., 2010). Similarly, Fortune et al. reported correlations between optic nerve rim parameters and axon count of r = 0.70 to 0.90 in experimental monkey glaucoma (Fortune et al., 2016). Our GliaR-visual function correlations fall within this range, supporting the validity of glial coverage metrics as structure-function indicators.

The biological plausibility of glial coverage area as a correlate of visual function rests on the established role of astrocytes in optic nerve pathophysiology. Reactive astrocytosis is among the earliest responses to glaucomatous injury, preceding overt axon loss in both D2 mice and induced ocular hypertension models (Son et al., 2010; Cooper et al., 2016). Bosco and colleagues demonstrated that glial coverage expands in direct proportion to axon loss across the disease spectrum (Bosco et al., 2016). Our glial coverage area ratio captures this phenomenon through automated segmentation, providing an objective metric that integrates information about both axon loss and reactive gliosis.

The age-dependent shifts in structure-function relationships observed in this study have implications for biomarker selection across disease stages. At early timepoints, morphometric features related to axon shape and size had the strongest correlations with visual function, potentially reflecting subclinical axonal stress. During mid-disease, glial coverage area ratio emerged as the dominant correlate of visual function. At late stages, axon diameter became the strongest predictor, possibly reflecting selective loss of larger axons that disproportionately contribute to visual function (Glovinsky et al., 1991). The emergence of axon diameter as the dominant correlate at 12 months may also reflect the biphasic nature of axonal pathology in glaucoma. During early and mid-stage disease, inflammatory processes and cellular stress can cause axonal swelling, producing heterogeneous diameter changes that obscure the relationship between axon caliber and function. As disease progresses, these acutely stressed axons either recover or degenerate, while surviving axons undergo atrophy. By 12 months, the inflammatory component has largely resolved, and reduced axon diameter more directly reflects cumulative axonal degeneration rather than transient swelling. This interpretation aligns with ultrastructural studies demonstrating that axonal swelling precedes frank degeneration in glaucomatous optic neuropathy (Howell et al., 2007). These dynamics suggest that optimal biomarkers may vary with disease stages.

The clear strain separation observed in principal component analysis also likely reflects distinct disease mechanisms rather than simply different points along a single pathological continuum. D2 mice develop pigmentary dispersion glaucoma secondary to mutations in *Gpnmb* and *Tyrp1*, which cause iris stromal atrophy and pigment liberation into the anterior chamber (Anderson et al., 2002). In contrast, BXD51 mice develop primary angle closure glaucoma with iris-corneal adherence through mechanisms independent of the *Gpnmb* and *Tyrp1* mutations (Velrajan et al., 2024). Despite these mechanistic differences, both strains demonstrated consistent relationships between glial coverage area ratio and visual function, suggesting that glial responses to axonal injury may represent a common pathway regardless of the initial insult. This convergence supports the generalizability of glial coverage metrics as structure-function biomarkers across glaucoma subtypes.

Several limitations of our study design should be acknowledged. The sample size (n = 6 per strain) limits statistical power for strain-specific subgroup analyses, and the smaller sample size within sex-stratified groups (n = 3 per sex per strain) further reduces power to detect sex effects. Although some sexual dimorphism was observed in B6, the absence of sexual dimorphism in diseased strains may reflect convergent pathological remodeling that obscures baseline differences. Histological assessment was performed only at 12 months, precluding direct temporal alignment between morphometric and clinical measurements. The glial coverage area ratio includes both glial cells and extracellular matrix, and cannot distinguish between reactive astrocytosis, microglial activation, and scar tissue formation without complementary immunohistochemical characterization. Additionally, axon counts obtained using light microscopic images underestimate true axon totals compared to transmission electron microscopy due to its inability to resolve smaller unmyelinated fibers. However, this limitation applies equally across all strains and is unlikely to bias between-strain comparisons.

In conclusion, glial coverage area ratio correlates with visual function as strongly as axon count across mouse strains with distinct optic nerve phenotypes, supporting inclusion of non-axon morphometric features in structure-function models. Age-dependent shifts in the relative importance of different morphometrics suggest that optimal biomarkers may vary across disease stages. These findings provide rationale for expanding quantitative optic nerve metrics beyond traditional axon counting.

## Abbreviations

RGC: retinal ganglion cells
VA: visual acuity
OS: left eye
UTHSC: University of Tennessee Health Science Center
IOP: intraocular pressure
nAx: axon count
AxDen: axon density
GliaR: glial coverage area ratio
Sol: mean solidity
AxDiam: mean axon diameter
MyArea: mean myelin area
AxMyArea: mean axon-myelin area.

## DATA AVAILABILITY

Data available upon reasonable request.

## Author Contributions

**BC:** Conceptualization, Methodology, Validation, Software, Formal analysis, Visualization, Writing - original draft, Writing - review & editing. **WW:** Methodology, Resources, Investigation, Writing - review & editing. **TJH:** Methodology, Resources, Investigation, Writing - review & editing. **XW**: Investigation, Writing - review & editing. **LG**: Investigation, Writing - review & editing. **MYK**: Visualization, Writing - review & editing. **MMJ:** Conceptualization, Supervision, Project administration, Funding acquisition, Writing - review & editing.

