## Supplementary Materials for "Visual Function Correlates More Strongly with Glial Coverage than Axon Count Across Multiple Mouse Strains"

**Supplementary Figure S1. Representative optic nerve segmentation workflow across three mouse strains at 12 months.** Each panel displays the same optic nerve (ON) cross-section processed through sequential stages of the AxonDeepSeg-based morphometric pipeline. **Top row of each panel:** Binary masks generated by the segmentation pipeline. **Bottom row of each panel:** Corresponding overlay visualizations on the original grayscale image. **Column labels:** Grayscale = original p-phenylenediamine (PPD)-stained bright-field image; Whole ON mask/overlay = total nerve boundary delineating cross-sectional area (red); Axon+Myelin mask/overlay = segmented axon and myelin sheath complexes (purple); Glial mask/overlay = parenchymal regions (blue). ON cross-sections from **(A)** C57BL/6J (B6), representing control optic nerve with densely packed axons and minimal parenchymal space, **(B)** BXD51, representing intermediate phenotype with preserved axon density, and **(C)** DBA/2J (D2) representing severe glaucoma phenotype with marked axon loss and expanded parenchymal/glial regions are shown. Progressive increase in glial mask area (white regions in top row, blue in bottom row) representing increased glial coverage area ratio accompanying axonal degeneration from control (A) to severely affected (C) nerves is observed.

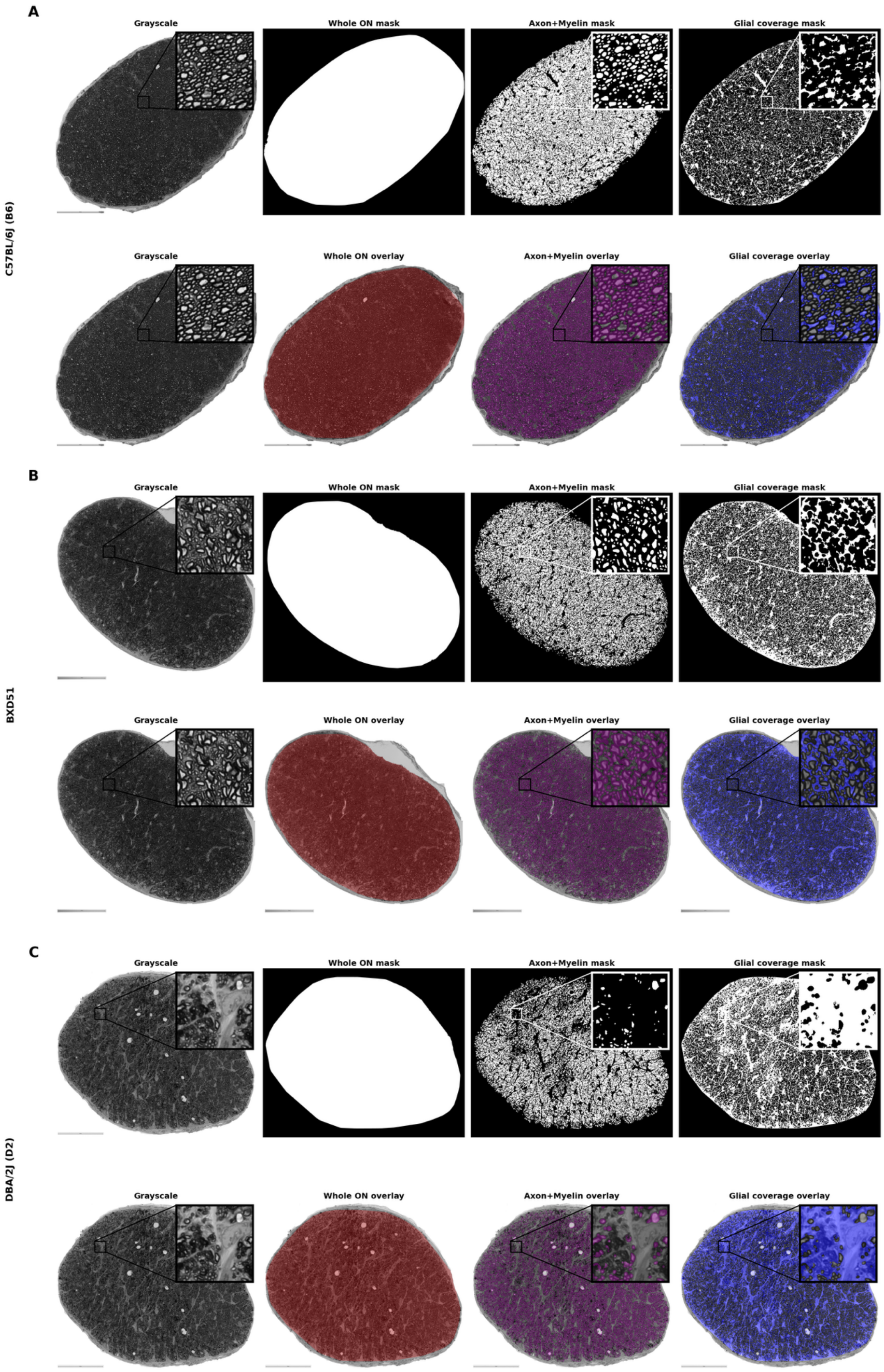
A
C57BL/6J (B6)
Grayscale
Whole ON mask
Axon+Myelin mask
Glial coverage mask
Grayscale
Whole ON overlay
Axon+Myelin overlay
Glial coverage overlay
B
BXD51
Grayscale
Whole ON mask
Axon+Myelin mask
Glial coverage mask
Grayscale
Whole ON overlay
Axon+Myelin overlay
Glial coverage overlay
C
DBA/2J (D2)
Grayscale
Whole ON mask
Axon+Myelin mask
Glial coverage mask
Grayscale
Whole ON overlay
Axon+Myelin overlay
Glial coverage overlay

**Supplementary Figure S2. Longitudinal clinical visual function profiles by strain.** Line plots display mean ± SD for four clinical measures across age (1, 3, 6, 9, and 12 months) in C57BL/6J (B6), BXD51, and DBA/2J (D2) mice from left eyes (OS) only (n = 6 per strain). **(A)** Visual acuity declines between 1-3 months in D2 mice, with values approaching zero by 9 months, whereas BXD51 maintains function until 12 months when values drop precipitously. B6 visual acuity declines between 1-3 months, then remains stable across ages. **(B)** Contrast threshold had sharp increase between 9-12 months in BXD51. **(C)** Intraocular pressure (IOP) is comparable across strains at early ages, with increases at 9-12 months most pronounced in BXD51. **(D)** Pattern electroretinography P50 (PERG P50) amplitude decreases across all stains by 12 months. **(E)** Absolute PERG N95 amplitude decreases across all strains by 12 months. Missing points indicate insufficient measurements at that time.

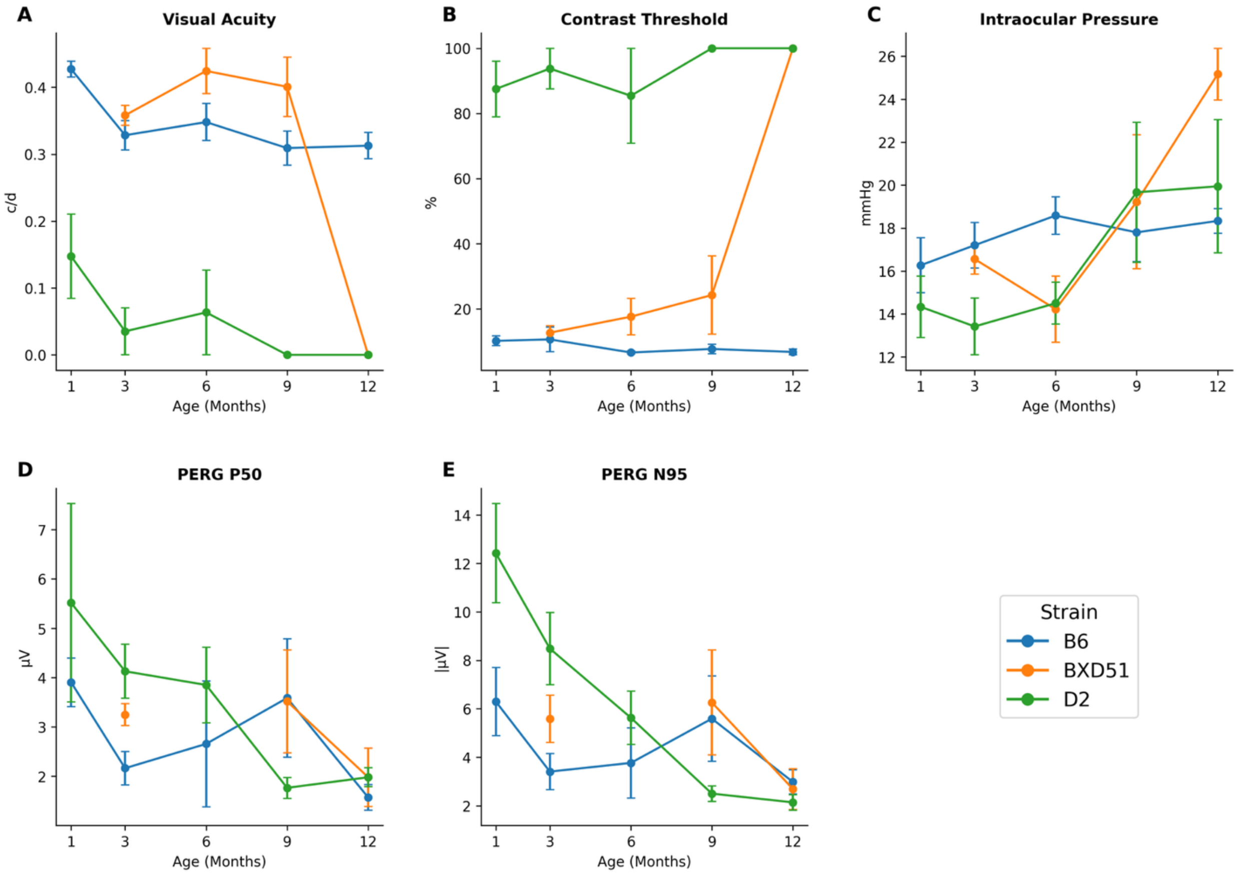

**Supplementary Figure S3. Sex-stratified optic nerve morphometrics by strain (median and IQR).** Box plots display median (center line) with interquartile range (IQR, box boundaries), with whiskers extending to 1.5 x IQR for six morphometric parameters in female and male mice across three C57BL/6J (B6), BXD51, and DBA/2J (D2) mice strains (n = 3 per sex per strain). Panels show **(A)** axon count, **(B)** glial coverage area ratio, **(C)** mean axon diameter, **(D)** mean myelin area, **(E)** mean axon+myelin area. **(F)** mean solidity.

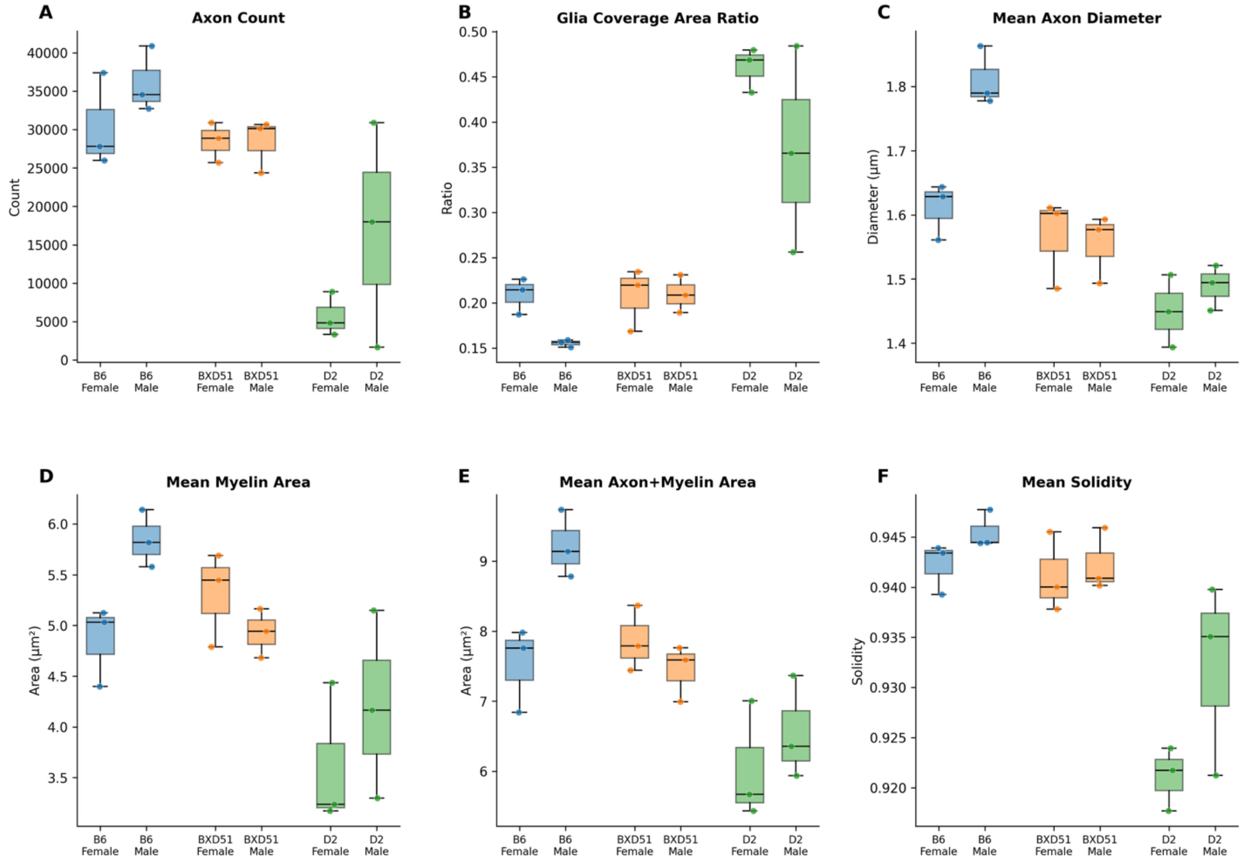

**Supplementary Figure S4. Sex-stratified optic nerve morphometrics by strain (mean and 95% confidence intervals). Bar graphs display mean values with 95% confidence intervals (error bars), with individual data points representing each mouse (n = 3 per strain). Six morphometric parameters** (A) axon count, (B) glial coverage area ratio, (C) mean axon diameter, (D) mean myelin area, (E) mean axon–myelin area, and (F) mean solidity **in female and male mice across** C57BL/6J (B6), BXD51, and DBA/2J (D2) mice strains are shown.

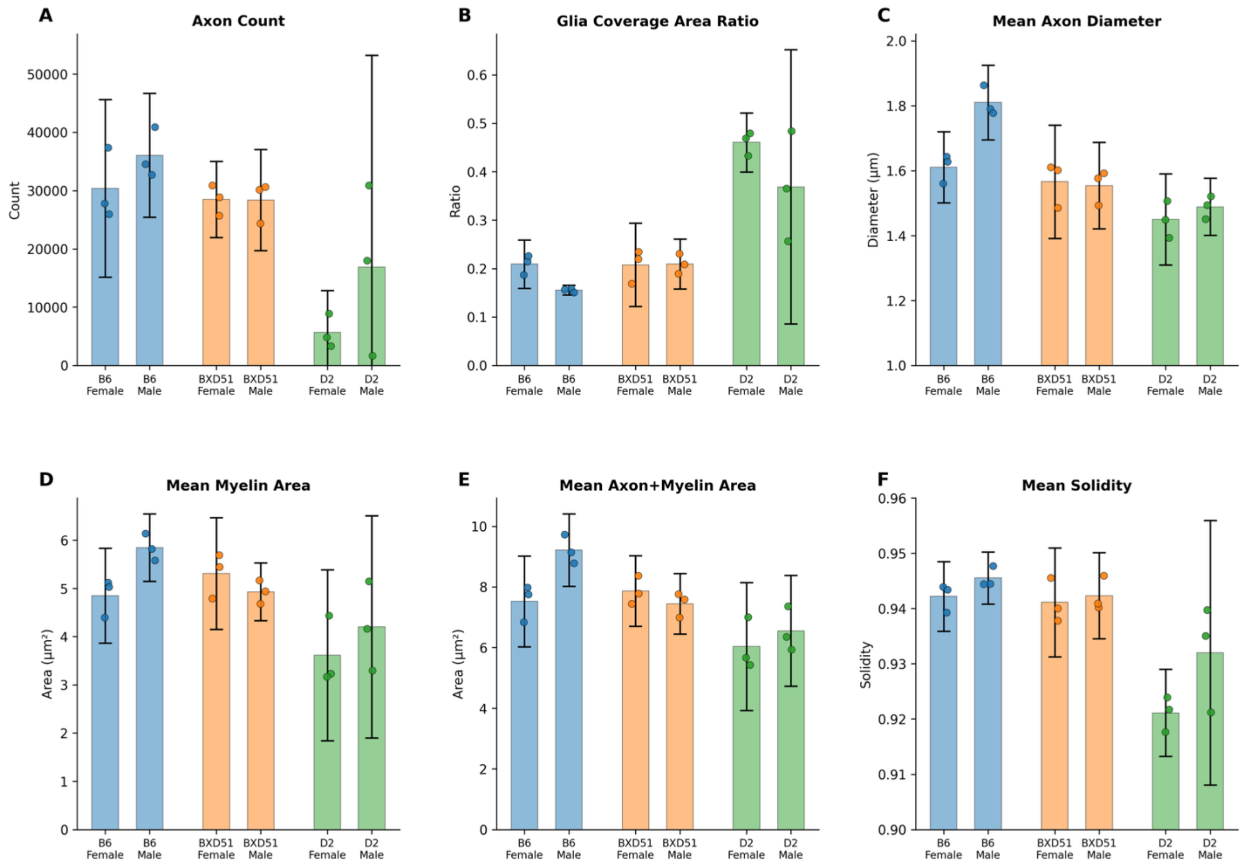

**Supplementary Table S1. Clinical Visual Function Profiles by Strain and Age**

| Clinical Measure | C57BL/6J (B6, n=6) | BXD51 (n=6) | DBA/2J (D2, n=6) | *p*-value |
|---|---|---|---|---|
| **Age: 1 Month** | | | | |
| Visual Acuity (c/d) | **0.43 ± 0.03** | **N/A** | **0.15 ± 0.15** | **.003**** |
| Contrast Threshold (%) | **10.15 ± 3.01** | **N/A** | **87.50 ± 20.92** | **<.001***** |
| IOP (mmHg) | 16.27 ± 2.86 | N/A | 14.33 ± 3.51 | .350 |
| PERG P50 (μV) | 3.91 ± 1.10 | N/A | 5.52 ± 2.84 | .285 |
| PERG N95 (μV) | -6.30 ± 3.14 | N/A | -12.42 ± 2.90 | .064 |
| **Age: 3 Months** | | | | |
| Visual Acuity (c/d) | **0.33 ± 0.05** | **0.36 ± 0.03** | **0.04 ± 0.09** | **<.001***** |
| Contrast Threshold (%) | **10.60 ± 8.53** | **12.66 ± 4.67** | **93.76 ± 13.95** | **<.001***** |
| IOP (mmHg) | 17.20 ± 2.36 | 16.56 ± 1.73 | 13.42 ± 2.64 | .057 |
| PERG P50 (μV) | **2.16 ± 0.76** | **3.25 ± 0.54** | **4.13 ± 1.10** | **.009**** |
| PERG N95 (μV) | **-3.41 ± 1.67** | **-5.58 ± 2.39** | **-8.48 ± 2.97** | **.024*** |
| **Age: 6 Months** | | | | |
| Visual Acuity (c/d) | **0.35 ± 0.06** | **0.42 ± 0.08** | **0.06 ± 0.16** | **<.001***** |
| Contrast Threshold (%) | **6.58 ± 0.82** | **17.58 ± 12.48** | **85.42 ± 35.72** | **<.001***** |
| IOP (mmHg) | 18.58 ± 1.75 | 14.22 ± 3.76 | 14.50 ± 2.38 | .073 |
| PERG P50 (μV) | 2.66 ± 2.55 | N/A | 3.85 ± 1.88 | .415 |
| PERG N95 (μV) | -3.77 ± 2.90 | N/A | -5.63 ± 2.69 | .327 |
| **Age: 9 Months** | | | | |
| Visual Acuity (c/d) | **0.31 ± 0.06** | **0.40 ± 0.10** | **0.00 ± 0.00** | **<.001***** |
| Contrast Threshold (%) | **7.66 ± 3.24** | **24.22 ± 26.87** | **100.00 ± 0.00** | **<.001***** |
| IOP (mmHg) | 17.80 ± 3.04 | 19.22 ± 7.64 | 19.67 ± 7.99 | .897 |
| PERG P50 (μV) | 3.59 ± 2.08 | 3.52 ± 2.56 | 1.77 ± 0.52 | .243 |
| PERG N95 (μV) | -5.59 ± 3.05 | -6.26 ± 5.31 | -2.50 ± 0.79 | .225 |
| **Age: 12 Months** | | | | |
| Visual Acuity (c/d) | **0.31 ± 0.04** | **0.00 ± 0.00** | **0.00 ± 0.00** | **<.001***** |
| Contrast Threshold (%) | **6.74 ± 1.98** | **100.00 ± 0.00** | **100.00 ± 0.00** | **<.001***** |
| IOP (mmHg) | 18.33 ± 1.00 | 25.17 ± 2.96 | 19.94 ± 7.61 | .150 |
| PERG P50 (μV) | 1.57 ± 0.58 | 1.98 ± 1.45 | 1.98 ± 0.47 | .733 |
| PERG N95 (μV) | -2.98 ± 1.13 | -2.69 ± 2.06 | -2.14 ± 0.79 | .626 |
| **Across All Ages (1, 3, 6, 9, 12 months)** | | | | |
| Visual Acuity (c/d) | **0.35 ± 0.06** | **0.30 ± 0.19** | **0.05 ± 0.11** | **<.001***** |

| | | | | |
|---|---|---|---|---|
| Contrast Threshold (%) | **8.34 ± 4.49** | **38.61 ± 39.11** | **93.32 ± 19.33** | **<.001*** ** |
| IOP (mmHg) | 17.53 ± 2.38 | 18.79 ± 5.97 | 16.58 ± 5.91 | .310 |
| PERG P50 (µV) | 2.71 ± 1.60 | 2.92 ± 1.76 | 3.05 ± 1.73 | .796 |
| PERG N95 (µV) | -4.33 ± 2.57 | -4.84 ± 3.71 | -5.01 ± 3.76 | .780 |

Data presented as mean ± SD for left eyes (OS) only. *P*-values from one-way ANOVA compare differences across three strains at each age (1, 3, 6, 9, and 12 months) and across all ages. N/A indicates data point not taken. Bold values indicate statistically significant strain differences (**p*<.05, ***p*<.01, ****p*<.001). *Abbreviations:* c/d = cycles per degree; IOP = intraocular pressure; PERG = pattern electroretinography.

**Supplementary Table S2. Sex-Stratified Morphometric Analysis**

| Strain | Sex | n | Axon Count | GliaR | Mean Axon Diameter (µm) | Mean Myelin Area ($\mu m^2$) | Mean Solidity |
|---|---|---|---|---|---|---|---|
| B6 | Female | 3 | 30,385 ± 6,130 | 0.209 ± 0.020 | 1.611 ± 0.044 | 4.851 ± 0.396 | 0.942 ± 0.003 |
| B6 | Male | 3 | 36,063 ± 4,279 | 0.155 ± 0.004 | 1.810 ± 0.046 | 5.846 ± 0.282 | 0.946 ± 0.002 |
| BXD51 | Female | 3 | 28,495 ± 2,633 | 0.208 ± 0.035 | 1.566 ± 0.070 | 5.309 ± 0.466 | 0.941 ± 0.004 |
| BXD51 | Male | 3 | 28,380 ± 3,486 | 0.210 ± 0.021 | 1.554 ± 0.054 | 4.930 ± 0.241 | 0.942 ± 0.003 |
| D2 | Female | 3 | 5,691 ± 2,877 | 0.460 ± 0.024 | 1.450 ± 0.056 | 3.614 ± 0.713 | 0.921 ± 0.003 |
| D2 | Male | 3 | 16,852 ± 14,652 | 0.369 ± 0.114 | 1.489 ± 0.035 | 4.203 ± 0.926 | 0.932 ± 0.010 |

**Two-Way ANOVA Results**

| Metric | Strain Effect (*p*-value) | Sex Effect (*p*-value) | Interaction (*p*-value) | B6 Sex Diff (t-test *p*) |
|---|---|---|---|---|
| Axon Count | **.0004*** ** | .119 | .410 | .259 |
| Glial coverage area ratio | **<.0001*** ** | .070 | .314 | **.010*** |
| Mean Axon Diameter | **<.0001*** ** | **.010*** | **.011*** | **.006*** ** |
| Mean Myelin Area | **.002*** ** | .154 | .134 | **.024*** |
| Mean Solidity | **<.0001*** ** | **.042*** | .226 | .143 |

| | | | | |
|---|---|---|---|---|
| Mean Axon+Myelin Area | **.0002*** ** | .060 | .035* | .019* |

Morphometric measurements are stratified by strain and sex as mean ± SD (n=3 per group). Two-way ANOVA tested main effects of strain, sex, and their interaction. Within-strain comparisons between males and females were performed using independent t-tests. Within-strain sex differences were observed in B6 only, with no differences detected in BXD51 or D2 (all *p*>.05). Bold values indicate statistically significant effects (**p*<.05, ***p*<.01, ****p*<.001). *Abbreviations:* GliaR = parenchymal area ratio.

**Supplementary Table S3. Pearson Correlations Between Morphometric Features and Clinical Measures by Age**

Age: 1 Month

| Clinical | Axon Count | Axon Density | GliaR | Solidity | Axon Diameter | Myelin Area | Axon-Myelin Area |
|---|---|---|---|---|---|---|---|
| **Visual Acuity** | **0.82** ** | **0.84** ** | **-0.86** ** | **0.86** ** | **0.75** ** | **0.85*** ** | **0.82** ** |
| **Contrast Threshold** | **-0.78** ** | **-0.81** ** | **0.85** ** | **-0.84** ** | **-0.86** ** | **-0.82** ** | **-0.86** ** |
| **Intraocular Pressure** | 0.59 | 0.47 | -0.49 | 0.53 | 0.39 | 0.37 | 0.36 |
| **PERG P50 Amplitude** | -0.66 | -0.64 | 0.66 | **-0.76*** | -0.32 | -0.48 | -0.36 |
| **PERG N95 Amplitude** | 0.20 | 0.20 | -0.39 | 0.39 | 0.63 | 0.43 | 0.54 |

Age: 3 Months

| Clinical | Axon Count | Axon Density | GliaR | Solidity | Axon Diameter | Myelin Area | Axon-Myelin Area |
|---|---|---|---|---|---|---|---|
| **Visual Acuity** | **0.84*** ** | **0.90*** ** | **-0.91*** ** | **0.86*** ** | **0.55*** | **0.79*** ** | **0.72** ** |
| **Contrast Threshold** | **-0.81*** ** | **-0.88*** ** | **0.92*** ** | **-0.85*** ** | **-0.65** ** | **-0.82*** ** | **-0.80*** ** |
| **Intraocular Pressure** | 0.19 | 0.29 | -0.34 | 0.25 | 0.33 | 0.28 | 0.32 |
| **PERG P50 Amplitude** | **-0.60*** | **-0.54*** | **0.66** ** | **-0.57*** | **-0.70** ** | **-0.66** ** | **-0.73** ** |
| **PERG N95 Amplitude** | 0.41 | 0.40 | **-0.54*** | 0.45 | **0.77** ** | 0.47 | **0.63*** |

Age: 6 Months

| Clinical | Axon Count | Axon Density | GliaR | Solidity | Axon Diameter | Myelin Area | Axon-Myelin Area |
|---|---|---|---|---|---|---|---|
| **Visual Acuity** | **0.91*** ** | **0.92*** ** | **-0.92*** ** | **0.88*** ** | **0.54*** | **0.76** ** | **0.71** ** |
| **Contrast Threshold** | **-0.93*** ** | **-0.93*** ** | **0.96*** ** | **-0.91*** ** | **-0.69** ** | **-0.82*** ** | **-0.80*** ** |
| **Intraocular Pressure** | 0.29 | 0.27 | -0.33 | 0.33 | **0.50*** | 0.36 | 0.41 |
| **PERG P50 Amplitude** | -0.10 | -0.29 | 0.29 | -0.23 | -0.30 | -0.27 | -0.30 |
| **PERG N95 Amplitude** | -0.04 | 0.15 | -0.16 | 0.08 | 0.26 | 0.11 | 0.19 |

Age: 9 Months

| Clinical | Axon Count | Axon Density | GliaR | Solidity | Axon Diameter | Myelin Area | Axon-Myelin Area |
|---|---|---|---|---|---|---|---|
| **Visual Acuity** | **0.76** ** | **0.81*** ** | **-0.83*** ** | **0.77*** ** | **0.53*** | **0.65** ** | **0.63** ** |
| **Contrast Threshold** | **-0.78*** ** | **-0.83*** ** | **0.85*** ** | **-0.8*** ** | **-0.64** ** | **-0.62*** | **-0.64** ** |
| **Intraocular Pressure** | **-0.49*** | -0.43 | 0.35 | -0.41 | -0.05 | -0.23 | -0.14 |
| **PERG P50 Amplitude** | 0.25 | 0.34 | -0.35 | 0.21 | 0.18 | 0.33 | 0.31 |
| **PERG N95 Amplitude** | -0.28 | -0.38 | 0.36 | -0.26 | -0.08 | -0.32 | -0.26 |

Age: 12 Months

| Clinical | Axon Count | Axon Density | GliaR | Solidity | Axon Diameter | Myelin Area | Axon-Myelin Area |
|---|---|---|---|---|---|---|---|
| **Visual Acuity** | **0.54*** | 0.43 | **-0.53*** | 0.46 | **0.77*** ** | 0.49 | **0.62*** |
| **Contrast Threshold** | **-0.52*** | -0.45 | **0.53*** | -0.47 | **-0.74** ** | -0.46 | **-0.58*** |
| **Intraocular Pressure** | 0.23 | 0.37 | -0.29 | 0.37 | -0.01 | 0.26 | 0.16 |
| **PERG P50 Amplitude** | 0.01 | -0.07 | 0.06 | 0.02 | -0.01 | -0.06 | -0.05 |
| **PERG N95 Amplitude** | -0.39 | -0.27 | 0.28 | -0.31 | -0.20 | -0.17 | -0.18 |

Values shown are Pearson correlation coefficients (r) between morphometric features and clinical visual function measures. Bold values indicate statistically significant correlations (**p*<.05, ***p*<.01, ****p*<.001). *Abbreviations:* GliaR = glial coverage area ratio; PERG = pattern electroretinography.
